# EXC-4/CLIC, Gα, and Rho/Rac signaling regulate tubulogenesis in *C. elegans*

**DOI:** 10.1101/2021.06.21.449267

**Authors:** Anthony F. Arena, Daniel D. Shaye

## Abstract

The Rho-family of small GTPases, which play crucial roles in development and disease, are regulated by many signal-transduction cascades, including G-protein-coupled receptor (GPCR)-heterotrimeric G-protein (Gα/β/γ) pathways. Using genetic approaches in *C. elegans* we identified a new role for Gα and Rho/Rac signaling in cell outgrowth during tubulogenesis and show that the Chloride Intracellular Channel (CLIC) protein EXC-4 is an evolutionarily-conserved player in this pathway. The gene *exc-4* was identified by its role in tubulogenesis of the excretory canal (*ExCa*) cell—a unicellular tube required for osmoregulation and fluid clearance. We identified an *exc-4* loss-of-function allele that affects an evolutionarily conserved residue in the C-terminus. Using this mutant we identified genetic interactions between *exc-4*, Gα, and Rho-family GTPases, defining novel roles for Gα-encoding genes (*gpa-12/*Gα_12/13_, *gpa-7/*Gα_i_, *egl-30*/Gα_q_, *gsa-1*/Gα_s_) and the Rho-family members *ced-10/Rac* and *mig-2/RhoG* in *ExCa* outgrowth. EXC-4 and human CLICs have conserved functions in tubulogenesis, and CLICs and Gα-Rho/Rac signaling regulate tubulogenesis during blood vessel development. Therefore, our work defines a primordial role for EXC-4/CLICs in Gα-Rho/Rac-signaling during tubulogenesis.

**One Sentence Summary:** Gα and Rho/Rac signaling regulates EXC-4/CLIC-mediated cell outgrowth during tubulogenesis in *C. elegans*, linking elements of G-protein signaling to the enigmatic CLIC family of proteins.

## INTRODUCTION

Biological tubes distribute essential nutrients throughout the body and collect the waste byproducts of cellular metabolism for excretion, making them indispensable for the development and viability of multicellular organisms. Tubulogenesis requires precise execution and coordination of cell behaviors (e.g., proliferation, migration, and outgrowth) often regulated by conserved signaling pathways (*1*). The *C. elegans* excretory system (Figure 1A) provides a simple yet powerful model to study the genetic control of tubulogenesis. This system is composed of three epithelial cells (the pore, duct, and <u>ex</u>cretory <u>ca</u>nal (*ExCa*) cell), which are connected to each other by inter-cellular adherens junctions (AJs), and each forms a unicellular tube during embryogenesis (reviewed in *2, 3*). The *ExCa* forms a tube by “hollowing”—a mechanism that also occurs during vertebrate angiogenesis (*4-6*). *ExCa* tubulogenesis initiates at the AJ that connects this cell to the duct, and lumen formation is associated with intracellular vesicles that appear to traffic towards this AJ, where they presumably fuse to form the nascent lumen (Fig. 1A, early embryo). A cytoskeletal structure, composed of F-actin, microtubules and intermediate filaments, assembles at the lumen-lining apical membrane to prevent swelling and promote lumen extension. Water flow into the growing lumen, via an apically-localized aquaporin, provides force for extension, and apically-directed vesicle trafficking and fusion provides membrane for lumen growth.

**Figure 1:**
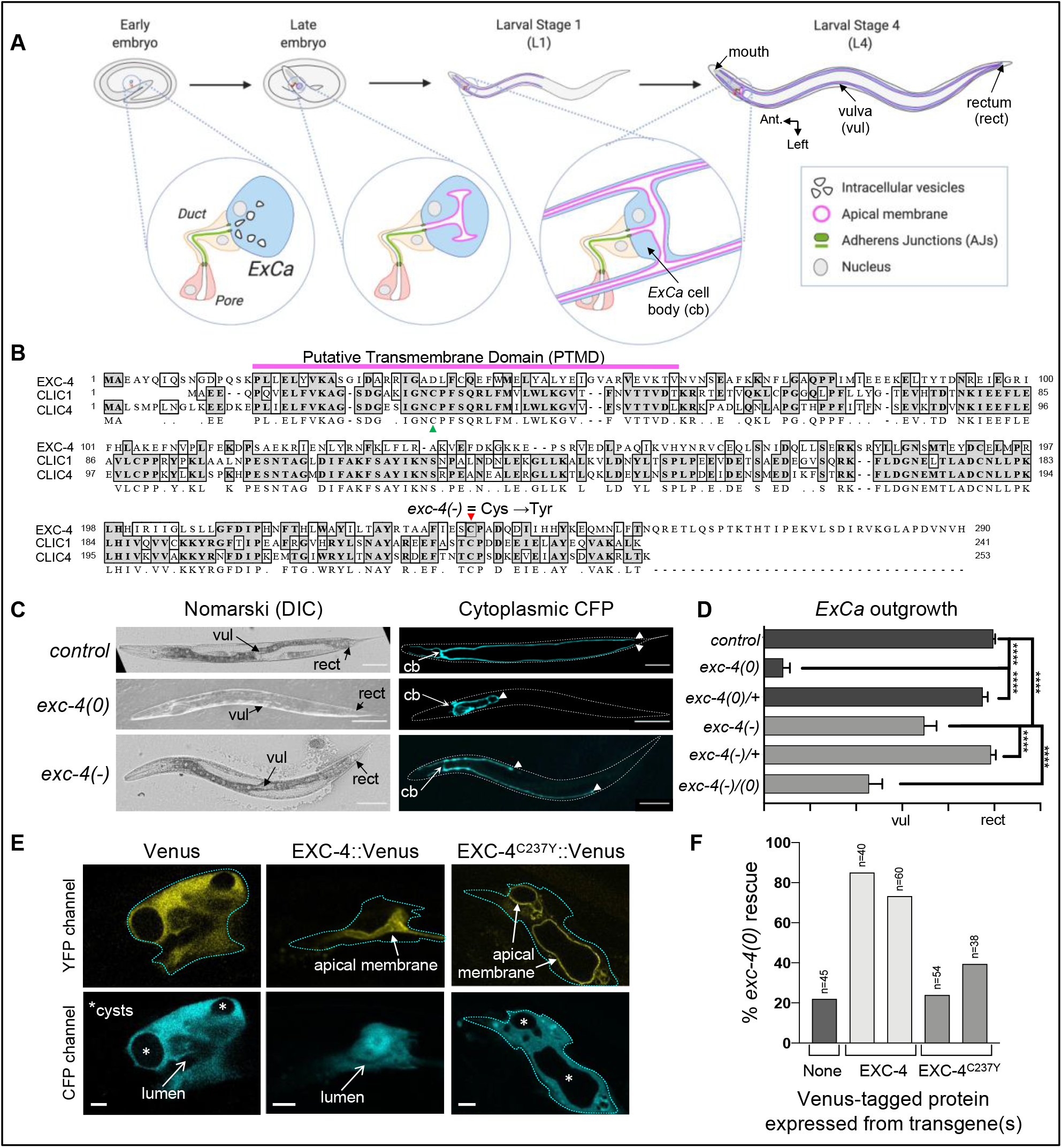
ExCa tubulogenesis and a new hypomorphic exc-4 allele. (**A**) the pore (red), duct (yellow), and *ExCa* (blue) cells form tubes during embryogenesis (reviewed in *2, 3*). These cells have a ring-shaped adherens junctions (AJs) connecting them to each other, and the duct has a longitudinal lumen-sealing AJ. In the early embryo vesicles in the *ExCa* appear to traffic towards the AJ, where they presumably fuse to form the nascent lumen. A microtubule, F-actin, and intermediate filament-rich cytoskeleton forms at the apical membrane (magenta) to restrict lumen diameter and promote extension. In late embryos the growing *ExCa* lumen bifurcates into left and right canals, and later these split once again into anterior and posterior canals. The lumen and *ExCa* basolateral membrane (dark blue line) grow out of the *ExCa* cell body (cb) in a coordinated manner. Posterior outgrowth is only partially done when animals hatch into L1 larvae, but completed before molting into the L2 stage. The L4 stage hermaphrodite depicted here highlights morphological features, like the vulva (vul) and rectum (rect), used to measure extent of posterior *ExCa* outgrowth (see Materials and Methods). **(B)** *C. elegans* EXC-4 aligned to human CLIC1 and CLIC4. The putative transmembrane domain (PTMD, magenta) is necessary and sufficient for EXC-4 apical membrane localization (*19, 24*). A cysteine required for GST-like activity (green) is not conserved in EXC-4. The cysteine mutated in *exc-4(-)* is conserved in all CLICs (red, and data not shown). (**C**) DIC and fluorescent images of control, *exc-4(0)*, and *exc-4(-)* hermaphrodites expressing cytoplasmic CFP (cyan) in the *ExCa*. Posterior *ExCa* tips (arrowheads) reach the rectum in controls, in *exc-4(0)* the *ExCa* forms large cysts and little outgrowth is seen, and in *exc-4(-)* the *ExCa* is not cystic, but posterior outgrowth is compromised. Scale bars here, and in Fig. 3, equal 100 µm. (**D**) quantification of posterior *ExCa* outgrowth shows that *exc-4(-)*is recessive and fails to complement *exc-4(0)*. **** = p≤0.0001. (**E**) confocal images of the *ExCa* cell body, marked with cytoplasmic CFP, in *exc-4(0)* hermaphrodites carrying transgenes expressing Venus-tagged proteins. Venus alone is cytoplasmic and does not rescue *exc-4(0)*. * = cysts caused by loss of *exc-4*. EXC-4::Venus rescues *exc-4(0)* and localizes to the lumen-lining apical membrane. Bright disk in this image is the *ExCa* nucleus, with dark nucleolus in the middle. EXC-4^C237Y^::Venus localizes to the apical membrane but fails to rescue *exc-4(0)*. Scale bars = 5 µm. (**F**) *exc-4(0)* rescue by EXC-4::Venus and EXC-4^C237Y^::Venus transgenes. Each bar represents an independent transgenic line. An outgrowth score >2 was set as the threshold for rescue. “None” controls are siblings of experimental animals with no detectable Venus in the *ExCa*.

As the lumen extends the *ExCa* basolateral membrane grows with it in a coordinated manner (*7*), a behavior termed “outgrowth”. *ExCa* outgrowth begins when this cell projects two hollow processes, or “canals”, from the left and right sides of the cell body. Later during embryogenesis these processes bifurcate into anterior and posterior canals that grow towards the head and tail—giving rise to this cell’s characteristic “H” shape (Fig. 1A, late embryo). When the embryo hatches into the first larval stage the anterior canals have completed outgrowth, while posterior canals have only grown ∼1/2 of their final length (Fig. 1A, L1 stage). Posterior outgrowth continues through the L1 stage (∼12 hours) until the posterior *ExCa* tips reach the rectum, and as the animal grows through three more larval stages (L4 stage in Fig. 1A) into adulthood the *ExCa* length to total body length ratio is maintained.

Genetic screens and candidate approaches have identified many conserved genes that regulate *ExCa* tubulogenesis (reviewed in *2, 3*) and several of these have been implicated in blood vessel tubulogenesis in vertebrates (*8-17*), demonstrating that these genetic approaches provide insights into conserved mechanisms of tubulogenesis. In addition, *ExCa* outgrowth and neuronal outgrowth share common features, such as guidance cues and cytoskeletal regulators, thus understanding the signaling pathways involved in *ExCa* outgrowth will also provide insight into neuronal guidance and outgrowth.

Here we focus on the gene *exc-4*, discovered in a screen for mutants with cystic *ExCa’s* (*18*), which encodes an ortholog of the conserved chloride intracellular channel (CLIC) family of proteins (Fig. 1B, and *19*). The discovery that EXC-4 regulates tubulogenesis in *C. elegans* suggested that CLICs regulate tubulogenesis in other contexts, for example during vascular development and angiogenesis. Indeed, CLIC1, CLIC4, and CLIC5 are expressed in distinct endothelial populations (*17, 20, 21*), CLIC1 and CLIC4 regulate angiogenic cell behaviors in culture (*14, 15*), and CLIC4 and CLIC5 regulate vascular development and function *in vivo* (*16, 17*), supporting the idea that CLIC proteins are conserved tubulogenesis regulators.

The first CLIC protein, p64/CLIC5B, was identified by its binding to the small molecule IAA94, a chloride channel inhibitor, and five more vertebrate CLICs have since been discovered and extensively studied (reviewed in *22*). Chloride channel activity for these proteins has not been unequivocally demonstrated *in vivo* or *in vitro* under physiological conditions, thus physiologically-relevant CLIC molecular function may not involve channel activity. In addition, structural studies have shown that CLICs resemble the Ω-class of glutathione-S-transferases (GSTs), and some CLICs have GST activity *in vitro*, but the physiological relevance of this activity also remains unknown. Notably, EXC-4 lacks a critical conserved cysteine required for GST function (Fig. 1B), thus the tubulogenesis-regulating function of EXC-4/CLIC does not appear to require GST-like function. CLICs have been linked to many cellular processes, including migration, survival, cytoskeletal regulation, differentiation, and intracellular trafficking. CLIC sub-cellular localization is as varied as the functions ascribed to this family, as they have been found in the cytoplasm, nucleus, at cell junctions, on intracellular vesicles, and at the plasma membrane (reviewed in *22*). In addition, CLICs can rapidly re-localize from one compartment to another in response to extracellular signals (*22, 23*). Given the plethora of molecular and cellular functions attributed to EXC-4/CLIC proteins, it is critical to define their *in vivo* function—a question we address here using *ExCa* tubulogenesis as a model.

Experiments aimed at assessing functional conservation between EXC-4 and CLIC1 revealed that they are interchangeable, but only when localized to the proper subcellular compartment. All CLICs have a putative transmembrane domain (PTMD) at their N-termini (Fig. 1B, and *22*). The EXC-4 PTMD drives constitutive apical plasma membrane localization in the *ExCa*, which is critical for EXC-4 function (*19, 24*). When human CLIC1 was expressed in the *ExCa* it was cytoplasmic and failed to rescue *exc-4* null, or *(0)*, mutants. However, when the CLIC1 PTMD was exchanged for the EXC-4 PTMD the chimeric protein localized at the apical membrane and efficiently rescued *exc-4(0)* phenotypes (*24*). Thus, although their PTMDs are functionally distinct, the EXC-4 and CLIC1 C-termini both promote tubulogenesis *in vivo*.

A potential role for CLICs in G-protein coupled receptor (GPCR)-heterotrimeric G-protein (Gα/β/γ)-Rho-family GTPase signaling was suggested by work showing that several GPCR ligands, including the potent angiogenic regulator sphingosine-1-phospate (S1P. Reviewed in *25*) regulate CLIC sub-cellular localization (*21, 26, 27*). S1P-receptor (S1PR) activation of G-protein coupled receptors (GPCRs), leading to dissociation of GPCR-bound heterotrimeric G-protein (Gα/β/γ) complexes and activation of downstream effectors, including the Rho-family GTPases RhoA, Rac1, and Cdc42 (*25, 28*). CLIC4 re-localization in response to S1P required Gσ13 and RhoA (*26*), suggesting that CLICs are regulated by Gα and Rho-family signaling. However, other studies, including recent work linking S1PR signaling to CLIC1 and CLIC4 function in endothelial cells (*26*), found that CLICs are required for RhoA and Rac1 activation (*21, 29, 30*), consistent with CLICs functioning upstream of Rho-family GTPases.

The results summarized above suggest that CLICs are regulators and/or effectors of the GPCR-Gα/β/γ-Rho-family pathway, but whether CLIC-mediated signaling regulates development and morphogenesis *in vivo* remained unknown. Using a new allele that changes a residue highly-conserved in CLIC proteins, we found that *exc-4* genetically interacts with Gα-encoding genes (*gpa-12/*Gα_12/13_, *gpa-7/*Gα_i_, *egl-30/*Gα_q_, *gsa-1/*Gα_s_), and with *ced-10/Rac* and *mig-2/RhoG*. Our work reveals a previously-unknown role for Gα and Rho/Rac signaling in *ExCa* outgrowth and shows that EXC-4/CLICs are evolutionarily conserved players in this signaling pathway.

## RESULTS

### A new hypomorphic allele of exc-4 identifies a specific role for CLICs in ExCa outgrowth

To ask if EXC-4/CLIC-mediated GPCR-Gα/β/γ-Rho/Rac signaling functions in *ExCa* tubulogenesis we wanted to test genetic interactions between *exc-4* and mutations in genes encoding GPCR-Gα/β/γ-Rho-family components. All previously-described *exc-4* alleles are nulls, resulting in a fully-penetrant cystic and shortened *ExCa* phenotype (Fig. 1C, D, and *18, 19*).

Because cysts in *exc-4(0)* mutants are very large, often spanning the entire internal cavity of the animal (Fig. 1C), and *ExCa* outgrowth is severely compromised (Fig. 1D. See Materials and Methods for outgrowth scoring criteria), it has not been possible to distinguish whether the *ExCa* outgrowth defect in *exc-4(0)* mutants reflects a specific role for EXC-4 in this process, or whether this phenotype is a secondary effect of cyst formation. The strength and penetrance of *exc-4(0)* alleles makes it challenging to assess whether mutations in other genes could enhance *exc-4* phenotypes. In addition, mutations in other genes that might suppress, or modify, *exc-4* activity may not be apparent if EXC-4 protein is completely absent and/or non-functional.

To identify potential hypomorphic (partial loss-of-function) *exc-4* alleles, which would provide a sensitized background for genetic interaction studies, we scanned the *C. elegans* “Million Mutation Project”, a library of mutagenized and whole-genome-sequenced strains (*31*), for mutations affecting conserved EXC-4/CLIC residues (Fig. 1B). We identified four candidate mutations and genetically characterized them (see Materials and Methods). Bona fide hypomorphic *exc-4* alleles should fit the following three criteria: 1) cause *ExCa* defects similar to, but milder than, *exc-4(0)*, 2) be recessive, and 3) fail to complement *exc-4(0)*. One mutation, changing a conserved C-terminal cysteine to tyrosine (Cys^237^ in EXC-4. Fig. 1B) fulfilled these criteria (Fig. 1C, D), and we refer to it as *exc-4(-)* hereafter. We found that *exc-4(-)* is temperature sensitive, as the outgrowth phenotype is more severe at higher temperatures (see Supp. Fig. 1)—a property we take advantage of in some of the genetic studies described below. In contrast to previously-characterized *exc-4(0)* mutants (*18, 19*), *exc-4(-)* animals do not form cysts (Figs. 1C), suggesting that the lumen maintenance and outgrowth functions of EXC-4 are separable and that Cys^237^ is required for the latter.

EXC-4 localization to the apical plasma membrane is critical for its function (*19, 24*), so we asked if the C237Y mutation affects subcellular localization. C-terminal tagging with the yellow fluorescent protein Venus does not affect EXC-4 apical localization (Fig. 1E) or function (Fig. 1F). In contrast, even though EXC-4^C237Y^::Venus localized to the apical membrane (Fig. 1E), it did not efficiently rescue (Fig 1F). Therefore, the conserved Cys^237^ residue is not required for proper protein localization but is required for EXC-4/CLIC function.

### Genetic interactions reveal a role for Gα orthologs in ExCa outgrowth

Cell culture studies have suggested a connection between CLICs and GPCR-Gα-Rho/Rac signaling (*21, 26, 27*), but a role for this pathway in *ExCa* tubulogenesis has not been identified.

*C. elegans* encodes ∼1,500 GPCRs, with at least one representative from each of the five GPCR super-families present (*32, 33*). Due to the expansion of this receptor family, and divergence in protein-coding sequences, it is challenging to assign orthology between human and *C. elegans* GPCRs (*34, 35*), and there are no clear *C. elegans* homologs of the GPCRs shown to regulate CLICs in cell culture (*21, 26, 27*) to test for *ExCa* function. Thus, we focused on proteins functioning downstream of GPCRs that have been associated with CLICs in cultured cells—the Gα and Rho-family GTPases (*21, 26*).

*C. elegans* has twenty-one Gα-like proteins (*36-38*), eight of which have identifiable human orthologs (*34, 35, 37, 38*). Null alleles of most of the conserved Gα genes are viable, and we obtained these to analyze *ExCa* outgrowth (see Materials and Methods). However, null *egl-30/*Gα_q_, *gsa-1*/Gα_s_, and *gpa-16*/Gα_i_ mutants are lethal (*39-41*), and for these we analyzed strong loss-of-function and/or activated alleles (see Materials and Methods). Individual Gα mutants did not display any *ExCa* defects (Fig. 2A-E, and Supp. Fig 2), so we assessed genetic interactions between these mutants and *exc-4(-)*. Loss of the *goa-1/*Gα_o_ or of three out of four Gα_i_ orthologs did not modify *exc-4(-)* (Supp. Fig. 2). In contrast, mutations in one Gα_i_ and the sole Gα_12/13_, Gα_q_, and Gα_s_ orthologs all displayed genetic interactions with *exc-4* as described below. Our analysis of these interactions indicate that Gα-mediated signaling plays a role in *ExCa* outgrowth.

**Figure 2:**
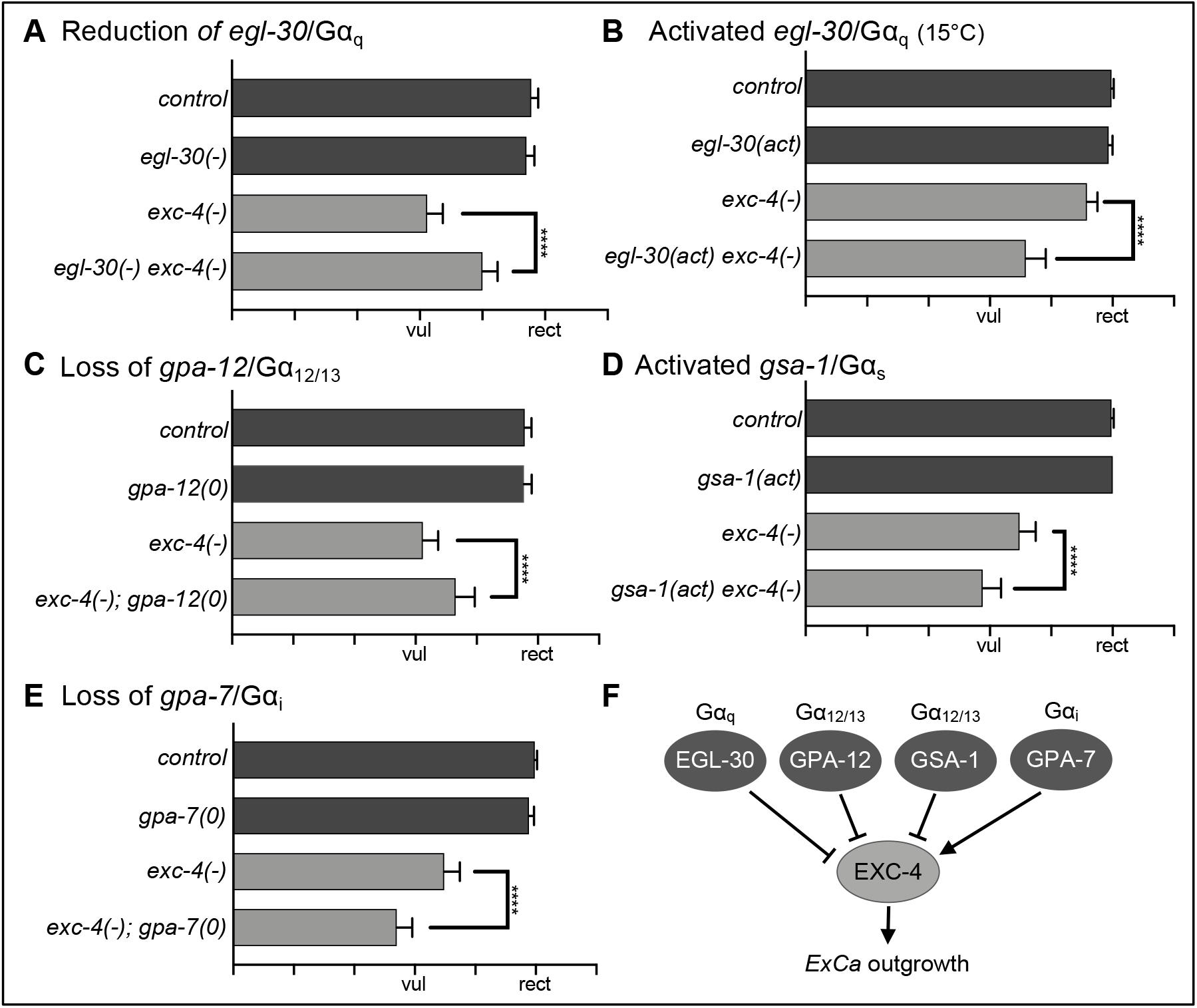
exc-4(-) genetically interacts with mutations in Gα-encoding genes. (**A**) Reduction of *egl-30*/Gσ_q_ function suppresses *exc-4(-)*, while an activated *egl-30* allele (**B**) shows the opposite genetic interaction. (**C**) A genomic deletion of *gpa-12*/Gσ_12/13_ suppresses *exc-4(-)*. (**D**) An activating mutation in *gsa-1*/Gσ_s_ and (**E**) loss of *gpa-7*/Gσ_i_ both enhance *exc-4(-)*. Based on these interactions, and results from cell culture studies suggesting that Gσ_12/13_ acts upstream of CLIC4 (*26*), we propose a model (**F**) for Gσ-mediated regulation of EXC-4 function in *ExCa* outgrowth.

A hypomorphic *egl-30/*Gα_q_ allele and a large deletion of *gpa-12*/Gα_12/13_ both significantly suppressed *exc-4(-)* (Fig. 2A, B). Based on studies in endothelial cells suggesting that Gα_12_ functions upstream of CLIC4 (26) (*26*), we propose that *egl-30/*Gα_q_ and *gpa-12/*Gα_12/13_ act as negative regulators of *exc-4/CLIC*, as their loss appears to counteract the reduced function of *exc-4(-)*. Notably, an *egl-30*-activating *(act*) mutation acted in the opposite manner, enhancing the weak *ExCa* outgrowth defect seen at 15°C (Fig. 2C and Supp. Fig. 1), supporting the idea that *egl-30/*Gα_q_ activity reduces *exc-4* function. We found activated *gsa-1*/Gα_s_ and a deletion of *gpa-7*/Gα_i_ significantly enhanced *exc-4(-)* outgrowth defects (Fig. 2D, E), suggesting that *gsa-1*/Gα_s_ is a negative regulator, while *gpa-7*/Gα_i_ is a positive regulator, of *exc-4* activity. Based on these observations we propose that each of these four Gα proteins modulates *exc-4* activity (Fig. 2F).The genetic interactions we observed also demonstrate that a functional interaction between Gα proteins and CLICs (*26*) is evolutionarily conserved

### The Rac ortholog ced-10 promotes ExCa outgrowth cell autonomously

Because Rho-family GTPases often function downstream of GPCR-Gα signaling (*42*), and because cell culture studies have shown that some Rho-family GTPases regulate (*26*), or are regulated by (*21, 30*), CLICs, we wanted to assess the function of Rho and Rac orthologs in *ExCa* outgrowth (*28, 42*). In addition, Rho-family GTPases are well-established regulators of neuronal outgrowth (*43-45*), and given that there are many players and mechanisms shared between *ExCa* and neuronal outgrowth in *C. elegans*, we hypothesized that Rho-family signaling may play a heretofore unrecognized role in *exc-4*-regulated *ExCa* outgrowth. The *C. elegans* genome encodes five canonical Rho-family GTPase orthologs: *rho-1/*Rho, *ced-10/*Rac, *rac-2*/Rac, *cdc-42/*Cdc42, and *mig-2/*RhoG (*46*). As previous studies have suggested that *rho-1* and *cdc-42* do not play major roles in *ExCa* outgrowth (*8, 47-50*) we focus here on potential roles for the Rac and RhoG orthologs *ced-10, rac-2*, and *mig-2* in *ExCa* outgrowth.

We analyzed three different hypomorphic *ced-10* loss-of-function mutations that disrupt different conserved regions (the C-terminal “CAAX” sequence required for CED-10/Rac membrane localization and the “Switch 1” or “Switch 2” regions that interact with upstream regulators and downstream effectors) and a large genomic deletion that is homozygous-sterile but maternally rescued (see Materials and Methods, Supplemental Fig. 3A, and *44, 45, 51*). These mutants all displayed significant *ExCa* outgrowth defects, with maternally-rescued *ced-10(0)* showing the strongest phenotype (Fig. 3A, B). A second *C. elegans* Rac, *rac-2*, is redundant with *ced-10* in some contexts (*44, 52, 53*), but not in others (*45, 54*). A *rac-2* deletion did not cause an *ExCa* phenotype on its own, nor did it exacerbate phenotypes caused by *ced-10* loss (Supp. Fig. 3B). Therefore, *rac-2* does not appear to function redundantly with *ced-10* in *ExCa* outgrowth.

**Figure 3:**
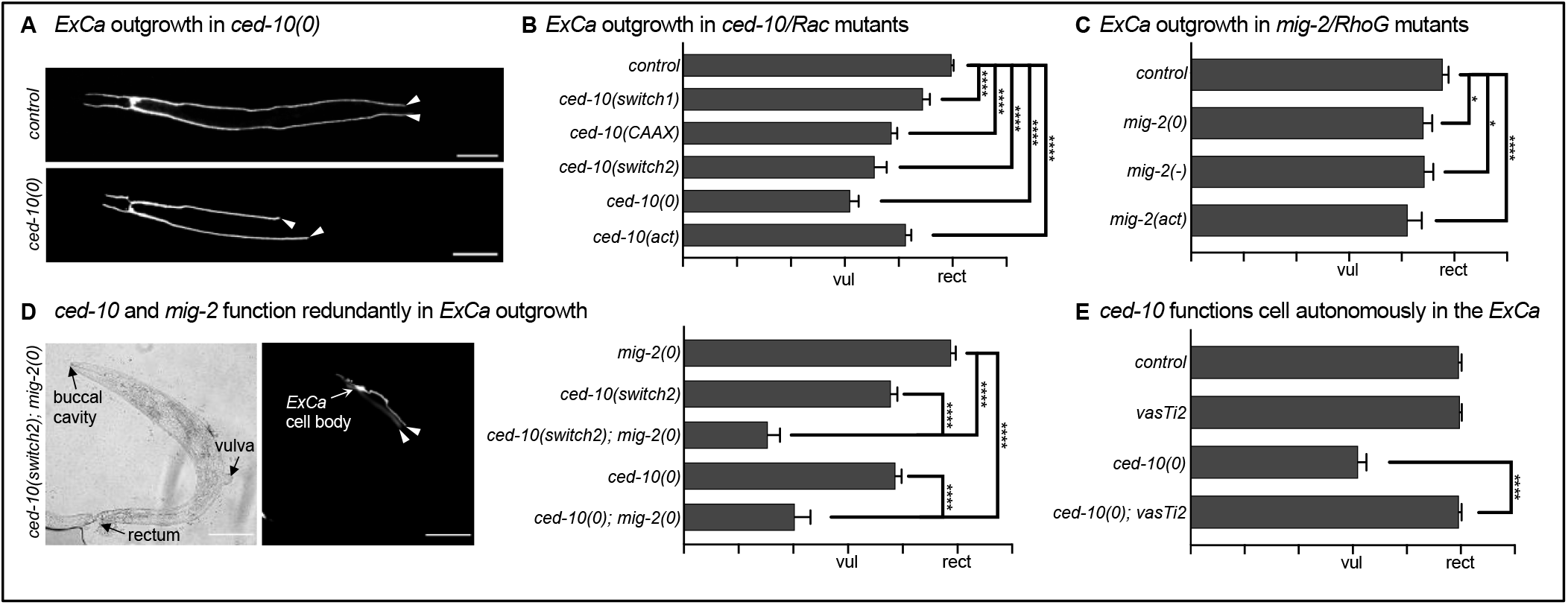
ced-10/Rac and mig-2/RhoG function redundantly to promote ExCa outgrowth. (**A**) fluorescent images of control and *ced-10(0)* hermaphrodites expressing cytoplasmic CFP in the *ExCa*. Arrowheads indicate extent of posterior *ExCa* outgrowth. (**B**) an allelic series of *ced-10/Rac* mutations (see Supp. Fig. 3A) cause significant outgrowth defects to varying extents. (**C**) different *mig-2/RhoG* mutations (see Supp. Fig. 3A) also cause significant outgrowth defects to varying degrees. (**D**) combining *ced-10* and *mig-2* loss-of-function mutations exacerbates *ExCa* outgrowth defects caused by each allele. (**E**) *vasTi2*, a single-copy genomic insertion transgene carrying FLAG-tagged *ced-10* under the control of the *ExCa* promoter *glt-3*p (see Supp. Fig. 3D) does not cause *ExCa* shortening on its own and significantly rescues *ExCa* outgrowth in *ced-10(0)*.

We tested a CRISPR-engineered *ced-10-*activating *(act*) mutation (Supplemental Fig. 3A and Ref. *45*) that, unexpectedly, did not act in a dominant manner (i.e., phenotypes were not seen in heterozygous animals. Supp. Fig. 3C) but caused significant *ExCa* shortening when homozygous (Fig. 3B). Genetic interactions between *exc-4(-)* and hypomorphic *ced-10* mutations were different than those seen between *exc-4(-)* and *ced-10(act)* (Fig. 4A, B), suggesting that the loss-of-function and activating mutation indeed have distinct effects on *ced-10* activity. The finding that loss-of-function and activating mutations cause similar phenotypes has been reported in studies of *ced-10/Rac* and *mig-2/RhoG* function in neurite outgrowth and cell migration (*43, 45*), and it has been suggested that this is likely related to the fact that GTPase function requires cycling between “on” and “off” states (*43*). We note that, similar to the *exc-4(-)* allele, *ced-10* mutations affect outgrowth but do not cause a cystic phenotype (Fig. 3A and data not shown), suggesting that *ced-10/Rac* functions specifically in *ExCa* outgrowth and not in lumen diameter maintenance.

**Figure 4:**
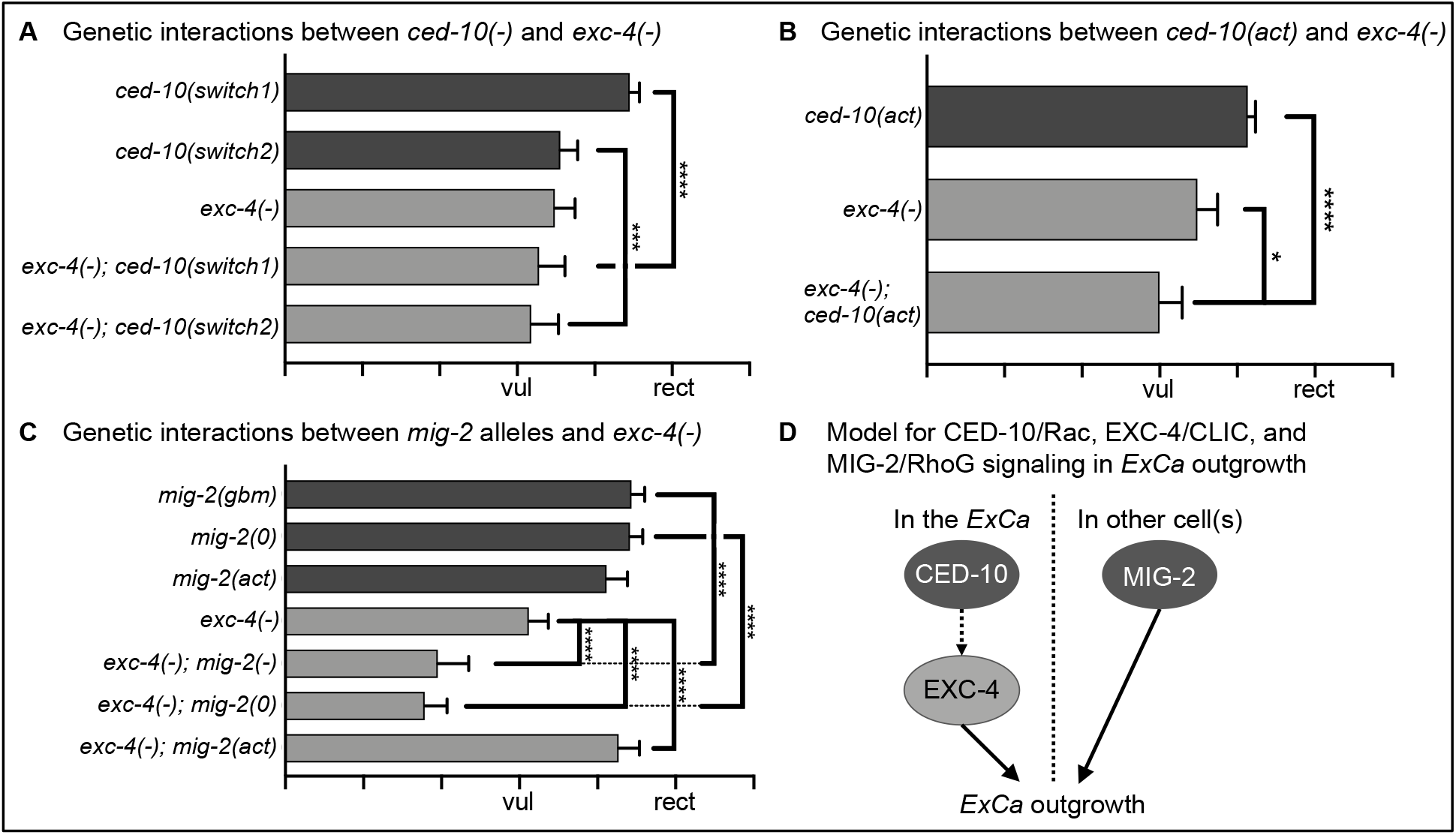
Genetic interactions between exc-4(-) and ced-10/Rac or mig-2/RhoG mutants. (**A**) two different *ced-10* loss-of-function alleles do not exacerbate *exc-4(-)*. (**B**) an activating mutation in *ced-10* does not suppress, but instead weakly enhances (*p=0.01), *exc-4(-)*. (**C**) mutations that reduce or eliminate *mig-2* enhance, and a mutation that activates *mig-2* suppresses, *exc-4(-)*. (**D**) based on the genetic interactions and transgene rescue results we propose a model where *ced-10* and *exc-4* function in the *ExCa*, and *mig-2* functions in other cell(s) to promote *ExCa* outgrowth.

To define the cellular focus of *ced-10* function in *ExCa* outgrowth we created a single-copy genome-inserted transgene, *vasTi2[glt-3*p*::FLAG::ced-10]* (see Materials and Methods, and Supp. Fig. 3D) that expresses N-terminal FLAG-tagged CED-10 under the control of the *ExCa* promoter *glt-3*p (*55*). We found that *vasTi2* rescued the outgrowth phenotype of *ced-10(0)* mutants (Fig. 3E), demonstrating that *ced-10/Rac* acts in the *ExCa* to promote outgrowth.

### The RhoG ortholog mig-2 functions in parallel to ced-10/Rac to promote ExCa outgrowth

MIG-2 has primary sequence similarity to Rac1 and Cdc42, but functions in a fashion similar to RhoG (*46, 56*). In *C. elegans ced-10/Rac* and *mig-2/RhoG* often function redundantly to regulate cell migration and outgrowth (*44, 45, 52, 53, 57, 58*). We analyzed several *mig-2* mutants (see Materials and Methods, and Supp. Fig. 3A): a hypomorphic allele affecting a guanine-binding motif, an early stop that acts as molecular null, and an activating mutation. These mutants all showed mild, but significant *ExCa* outgrowth defects (Fig. 3C). We built several combinations of *ced-10; mig-2* double mutants (See Materials and Methods) to assess functional redundancy, and saw a more severe *ExCa* outgrowth defect than in either single mutant (Fig. 3D, E), indicating that *ced-10/Rac* and *mig-2/RhoG* act redundantly in *ExCa* outgrowth. We note that, as for *ced-10* and *exc-4(-)* mutants, *mig-2* mutants affect outgrowth without causing a cystic phenotype (Fig. 3C, and data not shown), and even in the severe *ced-10; mig-2* doubles we did not observe *ExCa* cysts (Fig. 3D), supporting the idea that *ced-10/Rac* and *mig-2/RhoG* regulate *ExCa* outgrowth but not lumen diameter maintenance.

We asked if *mig-2*, like *ced-10*, functions cell autonomously to promote *ExCa* outgrowth. Because *mig-2(0)* animals exhibit only a mild outgrowth defect (Fig. 3B) we tested rescue in the more severe *ced-10(0); mig-2(0)* mutants (*44*). As for *ced-10*, we created a single-copy genomic-inserted transgene, *vasSi2[glt-3*p*::FLAG::mig-2]*, which was fully sequenced to validate its integrity (see Materials and Methods). However, unlike what we saw with *ced-10*, the *mig-2* transgene did not rescue *ExCa* outgrowth (Supp. Fig. 3E). Failure to rescue is unlikely due to a defect with the *glt-3*p promoter, as we continue to see strong *ExCa* expression from this promoter in *ced-10(0); mig-2(0)* doubles (Fig. 3D), or due to the FLAG tag, as N-terminally-tagged *mig-2* had been previously shown to be functional (*44*). Therefore, the failure to rescue the *ExCa* defect of *ced-10(0); mig-2(0)* doubles with *vasSi2[glt-3*p*::FLAG::mig-2]* suggests a cell non-autonomous role for *mig-2/RhoG* in *ExCa* outgrowth.

### ced-10/Rac and exc-4/CLIC function in the same pathway, which is parallel to mig-2/RhoG, to regulate ExCa outgrowth

Although combining *exc-4(0)* with *ced-10* or *mig-2* alleles resulted in sickness and/or lethality (data not shown). we were able to explore the functional relationship between *exc-4/CLIC, ced-10/Rac*, and *mig-2/RhoG* in *ExCa* outgrowth by assessing genetic interactions using different loss-of-function and/or activating mutations. As *exc-4* and *ced-10* both promote outgrowth cell autonomously (Fig. 3F and *19*), we hypothesized that they function in a linear pathway. We constructed *exc-4(-); ced-10(switch1)* and *exc-4(-); ced-10(switch2)* double mutants and found that neither of these doubles had a worse phenotype than the *exc-4(-)* single mutant (Fig. 4A), consistent with a model where *exc-4/CLIC* and *ced-10/Rac* function in the same pathway. We analyzed interactions between *exc-4(-)* and *ced-10(act)* to address the order of *exc-4* and *ced-10* function in *ExCa* outgrowth. If *ced-10/Rac* were activated by *exc-4/CLIC* then we would expect that the *exc-4(-)* defect might be suppressed by *ced-10(act)*. However, this was not the case (Fig. 4B), leading us to speculate that *exc-4* functions downstream of *ced-10* (Fig. 4D). We also note that *exc-4(-); ced-10(act)* doubles showed a slight, but significant, enhancement of the outgrowth defect seen in each single mutant (Fig. 4B). One interpretation of this result is that *ced-10(act)* reduces the compromised function of *exc-4(-)* even further, which would imply that *ced-10* acts as a negative regulator of *exc-4*.

If *ced-10* and *exc-4* function in the same pathway, then genetic interactions between *exc-4* and *mig-2* alleles should resemble those seen between *ced-10* and *mig-2* alleles (Fig. 3D, E). We constructed *exc-4(-); mig-2(-)* and *exc-4(-); mig-2(0*) strains, and these doubles showed a stronger phenotype than the respective single mutants (Fig. 4C), consistent with a model where *exc-4*, like *ced-10*, acts in parallel to *mig-2* in *ExCa* outgrowth. Combining *exc-4(-)* with an activating *mig-2* allele led to robust suppression of the *exc-4(-)* phenotype (Fig. 4C), indicating that activating *mig-2/RhoG* signaling can, at least partially, overcome a reduction in *exc-4/CLIC* function. In sum, our results reveal a new role for Gα proteins (which typically function upstream of Rho GTPases), *ced-10/Rac*, and *mig-2/RhoG* in *ExCa* tubulogenesis, and suggest a model where outgrowth is regulated cell autonomously by a *ced-10/Rac*-*exc-4/CLIC* pathway acting in parallel with *mig-2/RhoG* signaling, which likely occurs in a different tissue (Fig. 4D).

## DISCUSSION

The physiological and molecular functions of CLIC proteins have remained elusive over the last 30 years of research into this family of proteins. Recent work using cultured human cells implicate a link between GPCR-Gα-Rho/Rac signaling and CLIC function. Using the *ExCa* as model our work demonstrates that this functional interaction occurs *in vivo* and is evolutionarily conserved. CLICs have been found to function downstream of RhoA (*26, 59*), while other studies describe a function for CLICs upstream of RhoA and/or Rac1 (*21, 29, 30*). Our work indicates that that CED-10/Rac and EXC-4/CLIC act cell autonomously in the same pathway to regulate *ExCa* outgrowth (Fig. 4D). Our finding that a *ced-10* mutant with reduced GTPase activity, and thus locked in the active GTP-bound form (*45*), enhanced the *exc-4(-)* outgrowth defect led us to speculate that *ced-10* activity inhibits *exc-4* function. However, because mutations that affect Rho/Rac GTPase activity may also compromise wildtype function, it is challenging to deduce the exact effect of *ced-10/Rac* activity on *exc-4/CLIC* function based on these genetic interactions. Future genetic screens to identify mutations that suppress, or enhance, the *ExCa* outgrowth phenotype of *ced-10* and *exc-4* mutants or biochemical approaches to define CED-10 and EXC-4-interacting proteins should identify new players and illuminate how *ced-10/Rac* and *exc-4/CLIC* signaling are functionally coupled.

In addition to identifying a *ced-10/Rac-exc-4/CLIC* pathway that functions cell autonomously in *ExCa* outgrowth, we also found that *mig-2/RhoG* acts in parallel to this pathway in a cell non-autonomous manner. Previous work showed that the Rho/Rac guanine nucleotide exchange factor (GEF) *unc-73/Trio* had both cell autonomous and non-autonomous functions in *ExCa* guidance and outgrowth (*50*). Our results are consistent with this finding and suggest the possibility that the relevant *unc-73/Trio* targets are *ced-10/Rac* in the *ExCa* and *mig-2/RhoG* outside this cell. We speculate that a possible focus for *mig-2* function in *ExCa* outgrowth is the pair of canal-associated neurons (CANs), because their processes run in close proximity to the posterior *ExCa* arms along their length (reviewed in *2, 3*), and CAN positioning and outgrowth requires *mig-2* function cell autonomously.

Our work sheds light into cellular behaviors that underlie tubulogenesis. Functionally null *exc-4* alleles cause strong cystic and *ExCa* shortening phenotypes (*19*), but whether these phenotypes resulted from loss of separate *exc-4* functions remained unknown. In contrast, the new hypomorphic *exc-4* allele we report here only causes *ExCa* shortening, demonstrating that outgrowth and lumen maintenance are discrete *exc-4* functions. The fact that both *ced-10/Rac* and *mig-2/RhoG* mutants also display *ExCa* shortening, but not cystic phenotypes, also suggests that the cellular mechanisms underlying lumen maintenance and cell outgrowth are separable, with only the latter requiring Rho-family GTPases. We also found that the EXC-4^C237Y^ mutant protein encoded by *exc-4(-)* accumulates at the *ExCa* apical membrane, like wild-type EXC-4. This suggests that the conserved cysteine mutated in *exc-4(-)* is not required for proper sub-cellular localization and instead may be specifically involved in the signaling function of EXC-4/CLICs.

The conserved role for EXC-4/CLIC in Gα-Rho/Rac signaling we have defined here adds a new layer to the similarities between tubulogenesis in *C. elegans* and in endothelial cells. Recent work using human umbilical vein endothelial cells (HUVEC) showed that the angiogenesis-regulating lipid S1P, which activates GPCR-Gα/β/γ-RhoA/Rac1 signaling, induces rapid and transient re-localization of CLIC1 and CLIC4 from the cytoplasm to the plasma membrane, and that CLIC1 is required for S1P-induced RhoA and Rac1 activation, while CLIC4 only plays a role in Rac1 activation (*21*). Given that many critical players in GPCR-Gα/β/γ-RhoA/Rac1 signaling, including the Rho-family-regulating GEFs and GAPs (GTPase-activating proteins), are at least transiently localized to the plasma membrane and that this localization is critical for their function (*28, 60-62*), it is tempting to speculate that membrane-localized EXC-4/CLICs facilitate Rho/Rac signaling by promoting membrane accumulation of these players. Because EXC-4 is constitutively localized to the *ExCa* apical membrane, we suspect that membrane-bound EXC-4/CLIC-containing complexes may be more readily identified in the *ExCa* than in S1P-treated endothelial cells, where CLIC1 and CLIC4 membrane accumulation is transient (*21*). Thus, to further define EXC-4/CLIC function in Rho-family GTPase signaling we are focusing our efforts on identifying EXC-4-containing complexes in the *ExCa*. GPCRs, and the pathways regulated by these receptors, are such prevalent players in development and disease that ∼30% of all FDA-approved drugs target GPCR-mediated signaling (*63*). However, the pervasive role for GPCRs also makes it difficult to avoid harmful side effects from therapies that target their function. Therefore, the ability to define new players that couple

GPCRs to specific downstream effectors, like Gα proteins and Rho-family GTPases, will allow us to fully and safely harness the potential of targeting these pathways for therapy. Our work combined with findings in mammalian cells show that EXC-4/CLICs are components of membrane-bound complexes that regulate Rho-family signaling in different cellular and signaling contexts, suggesting that EXC-4/CLICs could be potential new targets for genetic and/or pharmacological manipulation of GPCR-Gα/β/γ-Rho family signaling.

## MATERIALS AND METHODS

### Strains

Unless otherwise noted, standard methods were used for strain handling and maintenance (*64*) and animals were grown at 20°C. Mutant alleles are described in WormBase (http://www.wormbase.org), and full genotypes of strains used in this study as well as primer sequences, plasmids created, and transgenes made from these plasmids are listed in Supplemental File S1. To identify *exc-4(-)* we scanned the Million Mutation Project (http://genome.sfu.ca/mmp/) for mutations that changed amino acids that are conserved between *C. elegans* EXC-4 and vertebrate CLICs. Four candidate mutations were identified and outcrossed ≥3x into a wildtype strain carrying the *ExCa-*marker *arIs198* (*55*) to remove background mutations and score *ExCa* phenotypes. Of the four candidates only *gk451333*, a missense mutation that changes Cys^237^ to Tyr, caused significant *ExCa* phenotypes and was characterized as described here.

### Imaging

Animals were mounted on 5% agar and immobilized with 2.5 mM levamisole followed by immediate imaging. For scoring outgrowth phenotypes animals were imaged on Zeiss AxioImager Z2 at 20x magnification. We visualized Venus and CFP from transgenes *vasSi1, vasEx1*, and *vasEx4* (Fig. 1E) using a Zeiss LSM880 confocal microscope at 40x magnification. Laser settings were: CFP (458 nm) laser power was 2.0, 2.6, and 2.0%, respectively for each *Ex* array. Venus (514 nm) power was 2.2, 2.0, and 2.2%, respectively, and TagRFP (561 nm) laser power was 1.0, 1.5, and 1.8%, respectively. Gain was kept equivalent for all images with CFP = 1000, Venus = 932, and TagRFP = 954. Confocal images acquired using AiryScan and post-processed in ImageJ using the same settings for all.

### Scoring ExCa outgrowth

The *ExCa* was visualized with transgenes expressing cytoplasmic Venus (*arIs164*) or cytoplasmic CFP and apically-localized LifeAct-TagRFP (*arIs198* or *arIs201*). Wildtype animals carrying these transgenes alone were used as controls. Each posterior *ExCa* arm grows independently of the other (*47, 55*), thus we scored each posterior canal in individual animals independently for outgrowth. Using morphological landmarks each posterior canal was scored on a scale from 0 – 5, where “0” indicates no outgrowth from the *ExCa* cell body, “3” represents outgrowth to the vulva and “5” corresponds to outgrowth to the rectum, and these landmarks are shown in the X axes in all graphs. For each genotype ≥55 *ExCa* posterior arms were scored. Bars in each graph denote the mean outgrowth score for the genotype shown and error bars represent the 95% CI. Significance was calculated by ANOVA with Sidak correction for multiple-comparisons using Prism software.

## ACKNOWLEDGMENTS

Early stages of this work were supported by Iva Greenwald (HHMI, Columbia University). We would like to thank Iva Greenwald, Jan Kitajewski, and Alexandra Socovich for critical reading of the manuscript. We are grateful to Jan Kitajewski, De Yu Mao, and Matt Kleinjan for helpful discussions, and we thank Julianna Escudero and Alison Kitajewski for technical assistance. We also thank Roger Pocock (Monash University) and the CGC, which is funded by NIH Office of Research Infrastructure Programs (P40 OD010440), for strains. This study was supported by an Research Supplement to Promote Diversity (NHLBI R01HL119403-02S1) and NIGMS R01 GM134032 to D.D.S, and an NSF GRFP (#1444315) to A. F. A.

## SUPPLEMENTARY MATERIALS AND METHODS

### Strains

Genotypes of all strains used are listed in the table below. Fluorescently-marked transgenes and balancers (*65, 66*) used for crosses, and to maintain sterile or lethal mutants, were:

Linkage Group (LG)1: *oxTi550,oxTi559, oxTi718, oxTi723, tmC20[Venus]*

LG3: *rhIs4*

LG4: *oxTi608, oxTi915, oxTi404, oxTi554, tmC25[Venus]*

LG5: *oxTi575, oxTi405*

LGX: *oxTi400, tmC24[mCherry]*

| Strain | Genotype | Shorthand | Figure(s) |
| --- | --- | --- | --- |
| GS6602 | *arIs198* (Supp. Ref. 55) | *control* | 1C, D; 2B, D, E; 3A, B<br>Supp. Figs 1, 2, 3B, C |
| GS8155 | *exc-4(rh133); arIs198* | *exc-4(0)* | 1C, D. |
| QN113 | *exc-4(gk451333); arIs198* | *exc-4(-)* | 1C, D; 2B, D, E; 4A, B<br>Supp. Figs 1, 2 |
| QN405 | *vasSi1 exc-4(rh133); arIs201* | *control* | 1E |
| QN373 | *exc-4(rh133); unc-119(ed3) pha-1(e2123); arIs198; vasEx1* | EXC-4::Venus | 1E, F |
| QN375 | *exc-4(rh133); unc-119(ed3) pha-1(e2123); arIs198; vasEx4* | EXC-4<sup>C237Y</sup>::Venus | 1E, F |
| QN374 | *exc-4(rh133); unc-119(ed3) pha-1(e2123); arIs198; vasEx9* | EXC-4::Venus | 1F |
| QN376 | *exc-4(rh133); unc-119(ed3) pha-1(e2123); arIs198; vasEx5* | EXC-4<sup>C237Y</sup>::Venus | 1F |
| QN244 | *arIs201* (Supp. Ref. 47) | *control* | 2A, C, and 3C |
| QN124 | *exc-4(gk451333); arIs201* | *exc-4(-)* | 2A, C, and 4C |
| QN138 | *egl-30(ad805); arIs201* | *egl-30(-)* | 2A |
| QN345 | *egl-30(ad805) exc-4(gk451333); arIs201* | *egl-30(-) exc-4(-)* | 2A |
| QN409 | *egl-30(ep271); arIs198* | *egl-30(act)* | 2B |
| QN410 | *egl-30(ep271) exc-4(gk451333); arIs198* | *egl-30(act) exc-4(-)* | 2B |
| QN135 | *arIs201; gpa-12(pk322)* | *gpa-12(0)* | 2C |
| QN283 | *exc-4(gk451333); arIs201; gpa-12(pk322)* | *exc-4(-); gpa-12(0)* | 2C |
| GS7951 | *gsa-1(ce81); arIs198* | *gsa-1(act)* | 2D |
| QN341 | *gsa-1(ce81) exc-4(gk451333); arIs198* | *gsa-1(act) exc-4(-)* | 2D |
| QN392 | *gpa-7(pk610); arIs198* | *gpa-7(0)* | 2E |
| QN412 | *exc-4(gk451333); gpa-7(pk610); arIs198* | *exc-4(-); gpa-7(0)* | 2E. |
| QN110 | *ced-10(n3417)/tmC25[Venus]; arIs198* | *ced-10(0)* | 3A, B |
| QN214 | *ced-10(rp100); arIs198* | *ced-10(switch1)* | 3B, 4A |
| QN46 | *ced-10(n3246); arIs198* | *ced-10(switch2)* | 3B, 4A, Supp. Fig. 3C |
| QN215 | *ced-10(knu268); arIs198* | *ced-10(act)* | 3B, 4B, Supp. Fig. 3B |
| QN45 | *ced-10(n1993); arIs198* | *ced-10(CAAX)* | 3B, Supp. Fig. 3C |
| QN49 | *arIs201; mig-2(mu28)* | *mig-2(0)* | 3C |
| QN50 | *arIs201; mig-2(gm38)* | *mig-2(-)* | 3C |
| QN51 | *arIs201; mig-2(gm103)* | *mig-2(act)* | 3C |
| QN447 | *ced-10(n3417)/tmC25[Venus]; arIs164* | *ced-10(0)* | 3D, E; Supp. Fig. 3E |
| QN322 | *ced-10(n3246)/tmC25 [Venus]; arIs164; mig-2(mu28)* | *ced-10(switch2); mig-2(0)* | 3D |
| QN323 | *ced-10(n3246); arIs164* | *ced-10(switch2)* | 3D |
| QN438 | *ced-10(n3417)/tmC25[Venus]; arIs164; mig-2(mu28)* | *ced-10(0); mig-2(0)* | 3D, Supp. Fig. 3E |
| QN68 | *arIs164; mig-2(mu28)* | *mig-2(0)* | 3D, Supp. Fig. 3E |
| QN358 | *arIs164 vasTi2* | *vasTi2* | 3E |
| QN355 | *ced-10(n3417)/tmC25[venus]; arIs164 vasTi2* | *ced-10(0); vasTi2* | 3E |
| GS6125 | *arIs164* (Supp. Ref. 55) | *control* | 3E, Supp. Fig. 3E |
| QN204 | *exc-4(gk451333); ced-10 (n3246); arIs198* | *exc-4(-); ced-10(switch2)* | 4A |
| QN218 | *exc-4(gk451333); ced-10(rp100); arIs198* | *exc-4(-); ced-10(switch1)* | 4A |
| QN219 | *exc-4(gk451333); ced-10(knu268); arIs198* | *exc-4(-); ced-10(act)* | 4B |
| QN126 | *exc-4(gk451333); arIs201; mig-2(mu28)* | *exc-4(-); mig-2(0)* | 4C |
| QN167 | *exc-4(gk451333); arIs201; mig-2(gm103)* | *exc-4(-); mig-2(act)* | 4C |
| QN248 | *exc-4(gk451333); arIs201; mig-2(gm38)* | *exc-4(-); mig-2(gbm)* | 4C |
| QN388 | *gpa-16 (it143); arIs201* | *gpa-16(0)* | Supp. Fig. 2A |
| QN393 | *gpa-16 (it143) exc-4(gk451333); arIs198* | *gpa-16(0) exc-4(-)* | Supp. Fig. 2A |
| QN403 | *gpa-4(pk381) arIs201* | *gpa-4(0)* | Supp. Fig. 2A |
| QN404 | *exc-4(gk451333); gpa-4(pk381) arIs201* | *exc-4(-); gpa-4(0)* | Supp. Fig. 2A |
| QN353 | *exc-4(gk451333); odr-3(n1605); arIs198* | *exc-4(-); odr-3(0)* | Supp. Fig. 2B |
| QN377 | *odr-3(n1605); arIs198* | *odr-3(0)* | Supp. Fig. 2B |
| QN143 | *goa-1(sa734); arIs198* | *goa-1(0)* | Supp. Fig. 2C |
| QN289 | *goa-1(sa734) exc-4(gk451333); arIs201* | *goa-1(0) exc-4(-)* | Supp. Fig. 2C |
| QN48 | *rac-2(ok326); arIs198* | *rac-2(0)* | Supp. Fig. 3C |
| QN86 | *ced-10(n1993) rac-2(ok326); arIs198* | *ced-10(CAAX) rac-2(0)* | Supp. Fig. 3C |
| QN87 | *ced-10(n3246) rac-2(ok326); arIs198* | *ced-10(switch2) rac-2(0)* | Supp. Fig. 3C |
| QN470 | *vasSi2; arIs164* | *vasSi2* | Supp. Fig. 3E |
| QN472 | *vasSi2; ced-10(n3417)/tmC25[Venus]; arIs164; mig-2(mu28)* | *vasSi2; ced-10(0); mig-2(0)* | Supp. Fig. 3E |

### Primers and Geneblocks

Primers used to confirm genotypes and to amplify cDNA’s are listed in the table below. Restriction sites (underlined) and start/stop codons (bold) are shown in cDNA primers.

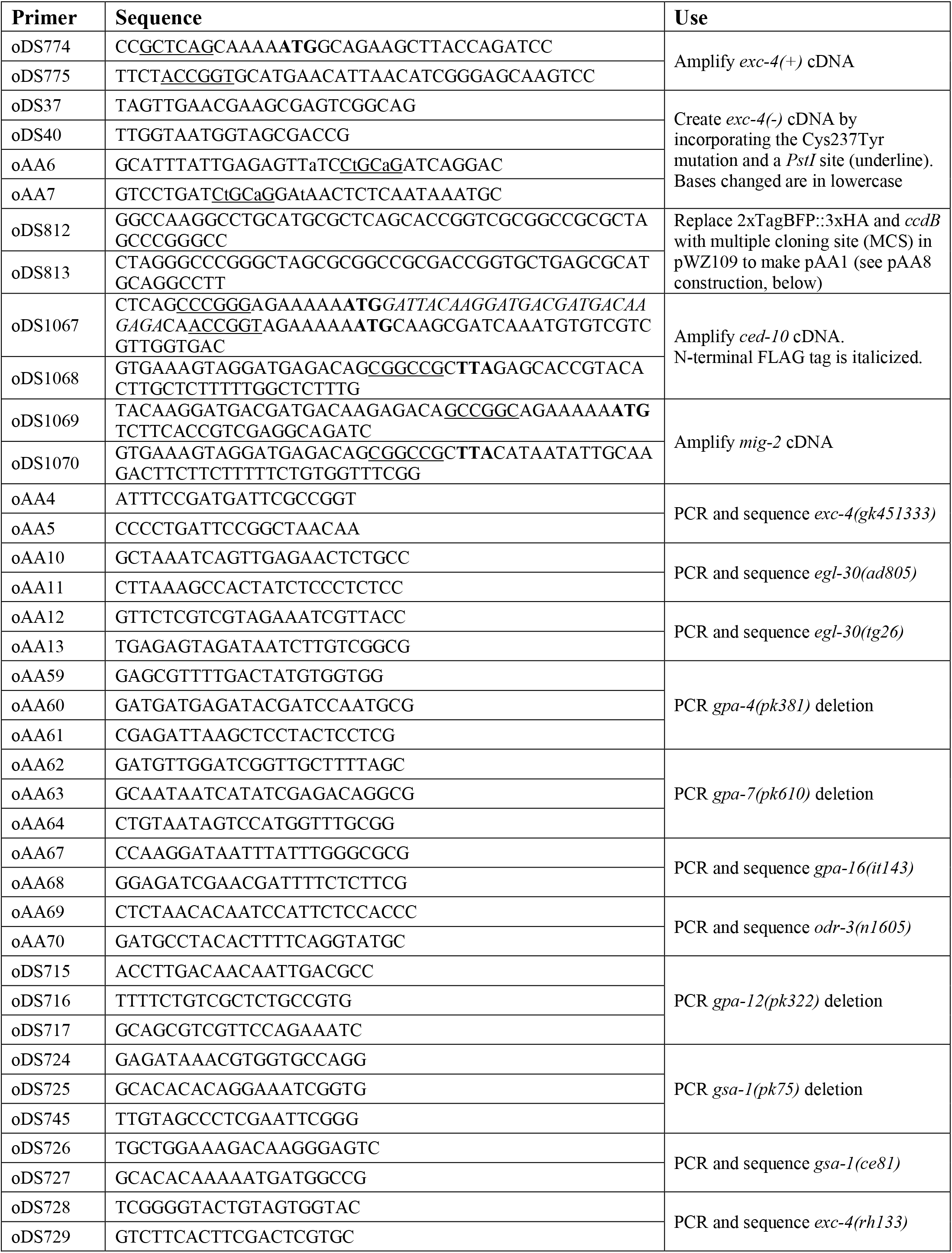
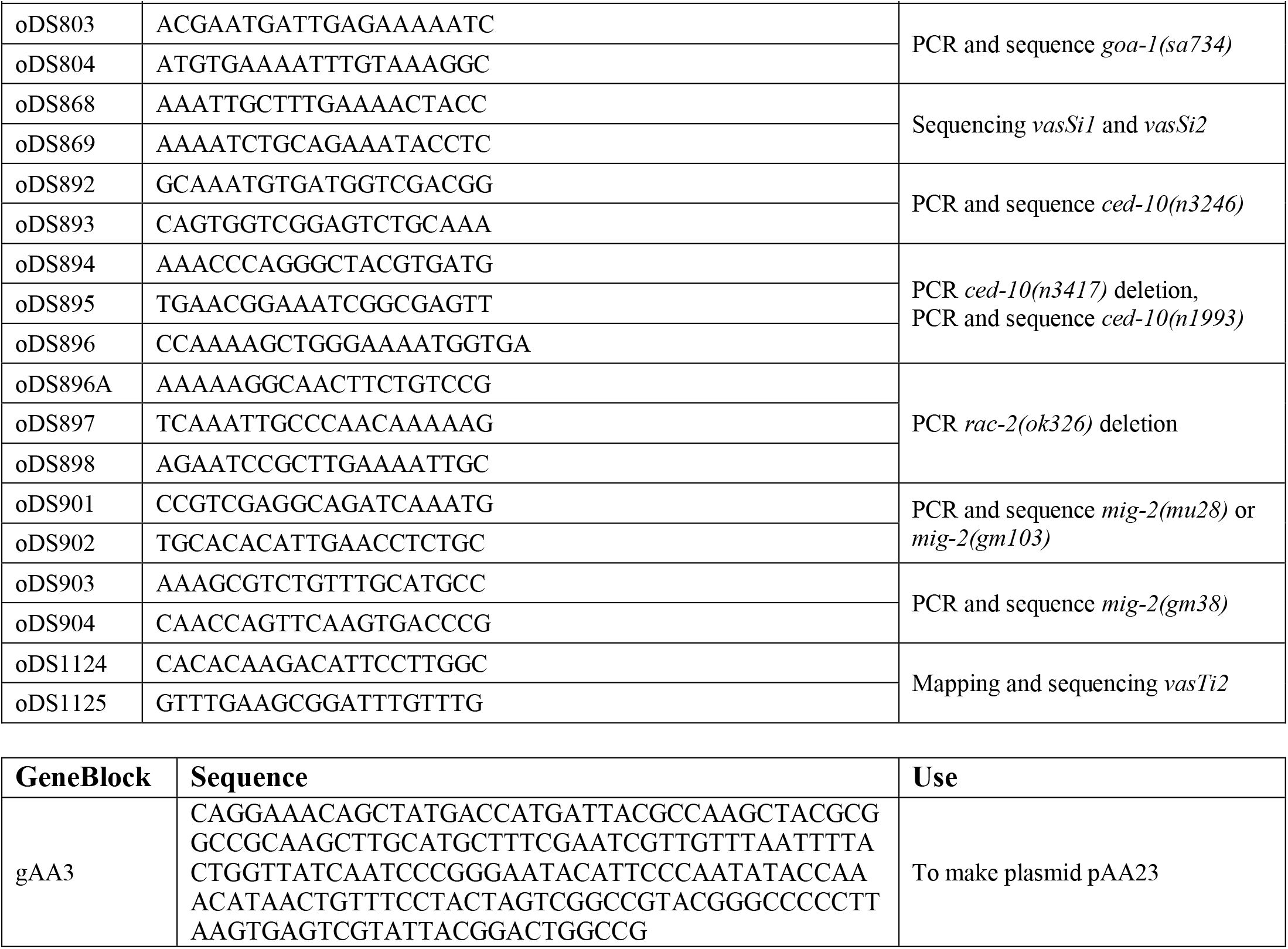

### Plasmids

#### pAA8 – *ttTi4348[glt-3*p*::Venus]*

Plasmid pWZ109 (a kind gift from A. Pani, D. Matus, and B. Goldstein) contains a self-excising cassette (SEC), for selection of CRISPR/Cas9-generated insertions (*67*), a TagBFP2 fluorescent reporter, and homology to the expression-permissive LG1 locus *ttTi4348* (*68*). The TagBFP2 portion of pWZ109 was removed by NotI/AvrII digestion and replaced with annealed and phosphorylated primers oDS812 and oDS813, creating a new plasmid, pAA1, which contains a multiple cloning site (MCS) and the SEC flanked by homology to *ttTi4348*. Plasmid pDS242, encoding the *Venus* flanked by the *ExCa* promoter *glt-3*p and the *unc-54 3’UTR* (*55*), was digested with SphI and ApaI and the *glt-*3p*::Venus::unc-54* 3’ fragment from this digestion was cloned into SphI/ApaI digested pAA1, generating pAA8. This plasmid was used to generate *vasSi1*, a single copy insertion in LG1 that expresses Venus in the *ExCa*, by CRISPR/Cas9 (*67*), as detailed in the “Transgenes” section below.

#### pAA14 – *glt-3*p*::exc-4::Venus*

The *exc-4* cDNA was previously PCR amplified and Gateway cloned into pDONR221 (Invitrogen) to make pDS323 (D. Shaye, unpublished). The *exc-4* cDNA from pDS323 was PCR amplified with oDS774 and oDS775. This PCR product was digested with AgeI/BlpI and cloned into AgeI/BlpI-digested pDS242 to create pAA14. pAA15 – *glt-3*p*::exc-4*^*C237Y*^*::Venus*. Using pAA14 as a template we performed two PCR reactions with primer pairs oDS37/oAA6 and oAA7/oDS40 to generate fragments with a 33bp overlap (bp 694-726 of *exc-4* cDNA). Primers oAA6 and oAA7 change the sequence ^709^TGT·CCC·GCC^717^, encoding Cys^237^-Pro-Ala, to TAT·CCT·GCA, encoding Tyr^237^-Pro-Ala and a *PstI* site. These fragments were used as templates in a second round of PCR, with primers oDS37 and oDS40, to obtain a “fused” fragment that was digested with AgeI and BlpI and ligated into AgeI/BlpI-digested pDS242, making pAA15.

#### pAA23 – Neo^R^-miniMos [glt-3p::Gibson::unc-54 3’UTR]

To avoid high levels of Rho GTPase expression, and possible associated pleiotropies, we wanted to make single-copy transgenes carrying *glt-3*p-driven *ced-10* and *mig-2* cDNAs. Plasmid pCFJ910 (a gift from Erik Jorgensen, Addgene plasmid # 44481), which contains a neomycin-resistant (Neo^R^) “mini” Mos1 transposon used to generate randomly-inserted single-copy transgenes (*66*), was digested with SnaBI and StuI, and the GeneBlock gAA3 (IDT) was introduced via Gibson assembly (*69*). The resulting plasmid, pAA23, encodes a *Neo*^*R*^-*miniMos* transposon with a unique restriction site (XmaI) flanked by 50 to 60 bp of homology to the 5’ end of the *glt-3* promoter and the 3’ end of the *unc-54* 3’UTR. Thus, we can excise various *glt-3*p-driven constructs from their original backbones and efficiently Gibson-clone them into XmaI-digested pAA23 to create *Neo*^*R*^*-miniMos* constructs to make random single-copy insertions.

#### pAA42 – Neo^R^-miniMos[glt-3p::FLAG::ced-10]

The *ced-10* cDNA was PCR amplified with primers oDS1067 and oDS1068, adding an N-terminal FLAG in frame to *ced-10*. This PCR product was digested with XmaI and NotI and inserted into AgeI/NotI-digested pDS242 to generate plasmid pJE13. This plasmid was then digested with SphI/SpeI to release the *glt-3*p*::FLAG::ced-10::unc-54 3’UTR* fragment, which was then Gibson cloned into XmaI-digested pAA23 to generate pAA42.

### pAA57 – Hygro^R^-ttTi4348[glt-3p::FLAG::mig-2]

One disadvantage of miniMos is that the single-copy transgenes insert randomly into the genome. As an alternative we used Mos1-mediated single-copy insertion (MosSCI), where a pre-existing Mos1 transposon in the genome is excised and replaced with the transgene of interest, which is flanked by homology to the genomic region neighboring the known excised Mos1 site (*68*). We made a hygromycin-resistant (Hygro^R^) MosSCI plasmid targeting the LG1 site *ttTi4348* by first digesting pAA1 (an intermediate created while making pAA8, see above) with KpnI/NheI to remove the large SEC. We then introduced the KpnI/SpeI fragment from pIR98 (*70*), which carries the Hygro^R^ cassette, making pAS100. To insert an *ExCa*-expressed *mig-2* cDNA into pAS100 we first amplified the *mig-2* cDNA with primers oDS1069 and oDS1070, digested the PCR product with XmaI/NotI and inserted this product into AgeI/NotI-digested pJE13 (see above), generating pJE16: a plasmid encoding *glt-3p::FLAG::mig-2::unc-54 3’ UTR*. Subsuqently pAS100 and pJE16 were SpeI/SphI digested and appropriate fragments were ligated to generate pAA57.

### Transgenes

All transgenes were made by germline micro-injection of different DNA constructs (*71*), as described below.

#### Extrachromosomal arrays *vasEx1, vasEx4, vasEx5* and *vasEx9*

*unc-119(ed3) pha-1(e2123ts); arIs198* hermaphrodites were injected with mixes containing plasmids of interest (see table below), and the co-injection markers pBX (*72*), which rescues *pha-1(e2123)*, and the Hygro^R^ plasmid pIR98 (*70*), both at 50ng/μl. Transformants were first selected by *pha-1* rescue at 22°C, thereafter arrays were followed by fluorescent expression from constructs, *pha-1* rescue, and/or Hygro^R^.

Single-copy transgene *vasSi1*[*glt-3p::Venus*]

plasmid pAA8, described above, was linearized with PciI and injected in a mix containing plasmid pCas9 (expresses Cas9) at 50 ng/µl, modified synthetic single-guide RNA (sgRNA) targeting the *ttTi4348* locus (Synthego) at 5 µM, and the co-injection markers pCFJ90 and pGH8 (2.5 and 5 ng/µl respectively) into wildtype (N2) hermaphrodites. After injection transformed animals and the resulting single-copy insertion were selected, and the SEC was excised, as previously described (*67*).

#### Single-copy transgenes *vasTi2*[*glt-3*p*::FLAG::ced-10*] and *vasSi2*[*glt-3*p*::FLAG::mig-2*]

protocols for mobilizing Mos1 (*66, 68*) were followed to make these transgenes. Mixes containing pAA42 or pAA57, described above, along with pCFJ90 (*myo-2::mCherry::unc-54 3’UTR*, 5ng/ μl), pMA122 (*hsp16*.*41::peel-1::tbb-2 3’UTR*, 20 ng/μl), and pCFJ60 (*eft-3::Mos1 transposase::tbb-2 3’ UTR*, 130 ng/μl), were injected into either wildtype (N2) hermaphrodites (for *vasTi2*) or into hermaphrodites carrying the *ttTi4348* Mos1 insertion (for *vasSi2*). Transformants were selected by *myo-2::mCherry* and growth on Neomycin (for pAA42) or Hygromycin (for pAA57). Heat-shock, which drives expression of the toxic protein PEEL-1, combined with the appropriate antibiotic medium were used to select against extrachromosomal arrays and identify the Neo^R^ miniMos insertion (*vasTi2*), or the Hygro^R^ replacement of the excised Mos1 in *ttTi4348* (*vasSi2*).

| Transgene | Construct | Plasmid | Concentration |
| --- | --- | --- | --- |
| *vasEx1* | *glt-3p::exc-4::Venus::unc-54 3'UTR* | pAA14 | 5 ng/μl |
| *vasEx9* |  |  |  |
| *vasEx4* | *glt-3p::exc-4(C237Y)::Venus::unc-54 3'UTR* | pAA15 | 5 ng/μl |
| *vasEx5* |  |  |  |
| *vasSi1* | *ttTi4348[glt-3p::Venus::unc-54 3'UTR]* | linearized pAA8 | 7 ng/μl |
| *vasTi2* | *miniMos[glt-3p::FLAG::ced-10::unc-54 3'UTR]* | pAA42 | 10 ng/μl |
| *vasSi2* | *ttTi4348 [glt-3p::FLAG::mig-2::unc-54 3'UTR]* | pAA57 | 50 ng/μl |

---

**Supp. Fig. 1:**
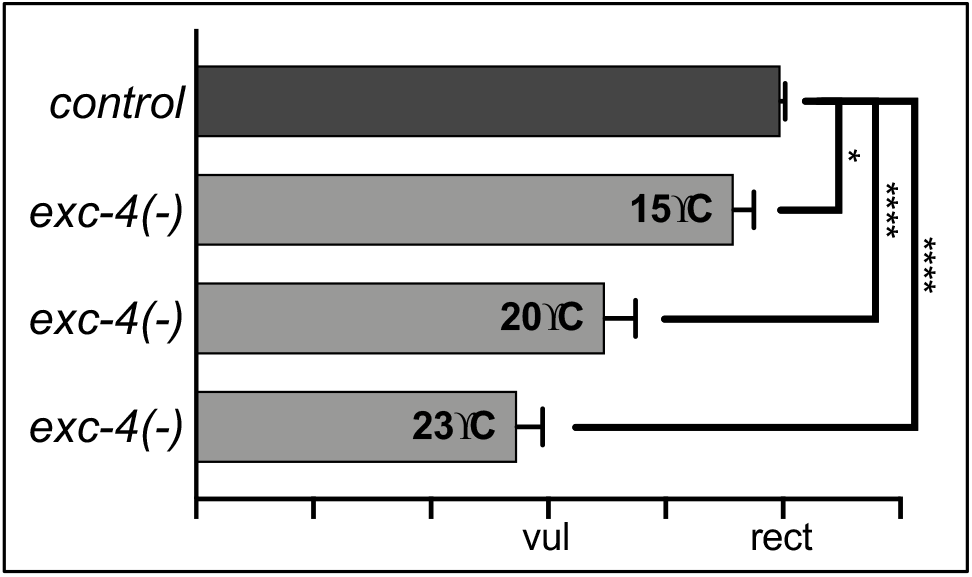
*exc-4(-*) is a temperature sensitive allele. *ExCa* outgrowth in *exc-4(-)* animals was measured, as described in Fig. 1B, at 15°C, 20°C, and 23°C, and phenotype severity increased with temperature.

**Supp. Fig. 2:**
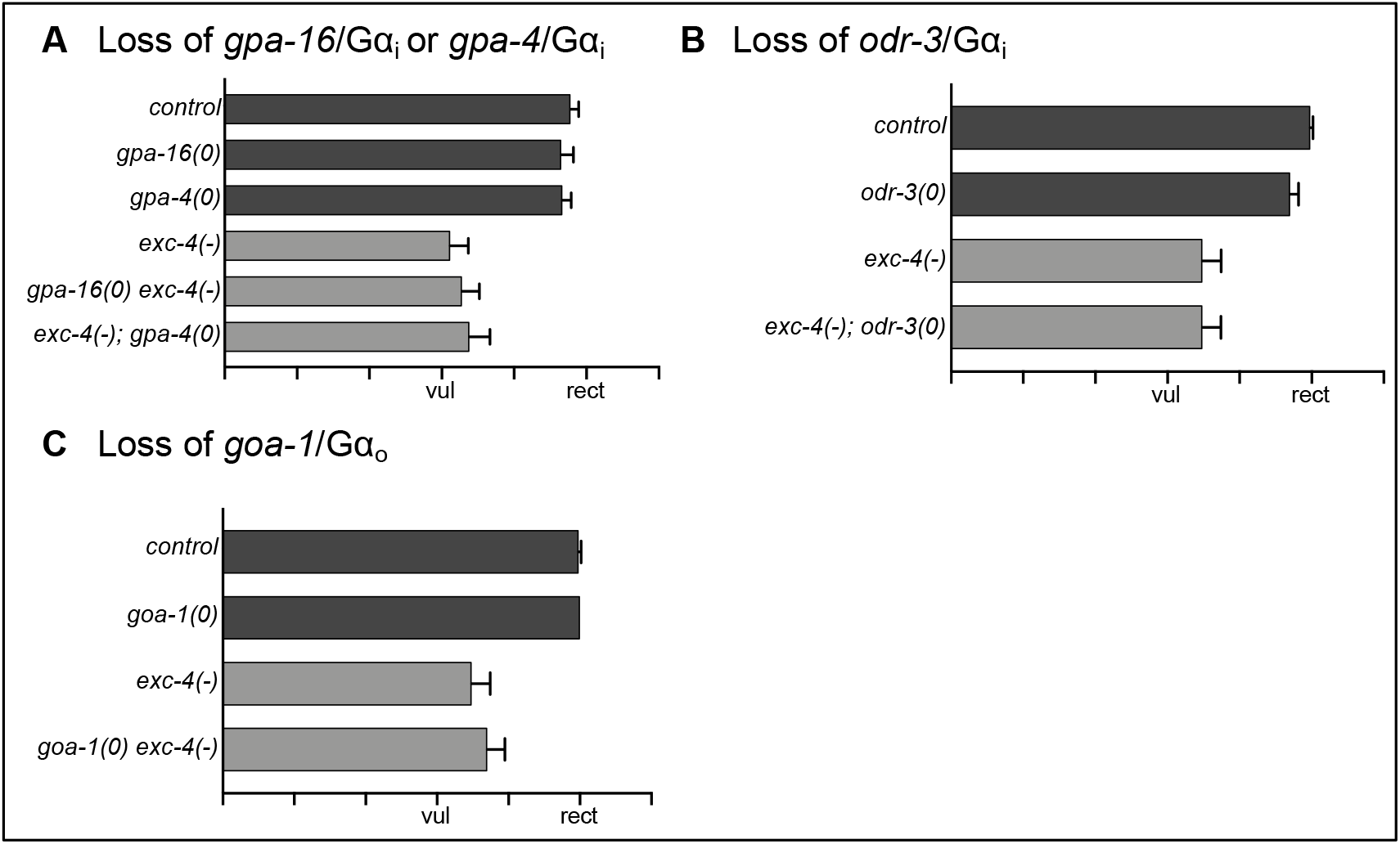
loss-of-function *mutations in three Gai paralogs, or in the sole Gao ortholog, do not cause ExCa outgrowth defects nor do they genetically interact with exc-4(-)*. Mutations in (**A**) *gpa-16*/Gσ_i_, *gpa-4*/Gσ_i_, (**B**) *odr-3/*Gσ_i_, or (**C**) *goa-1*/Gσ_o_ do not cause outgrowth defects on their own, or when combined with *exc-4(-)*.

**Supp. Fig. 3:**
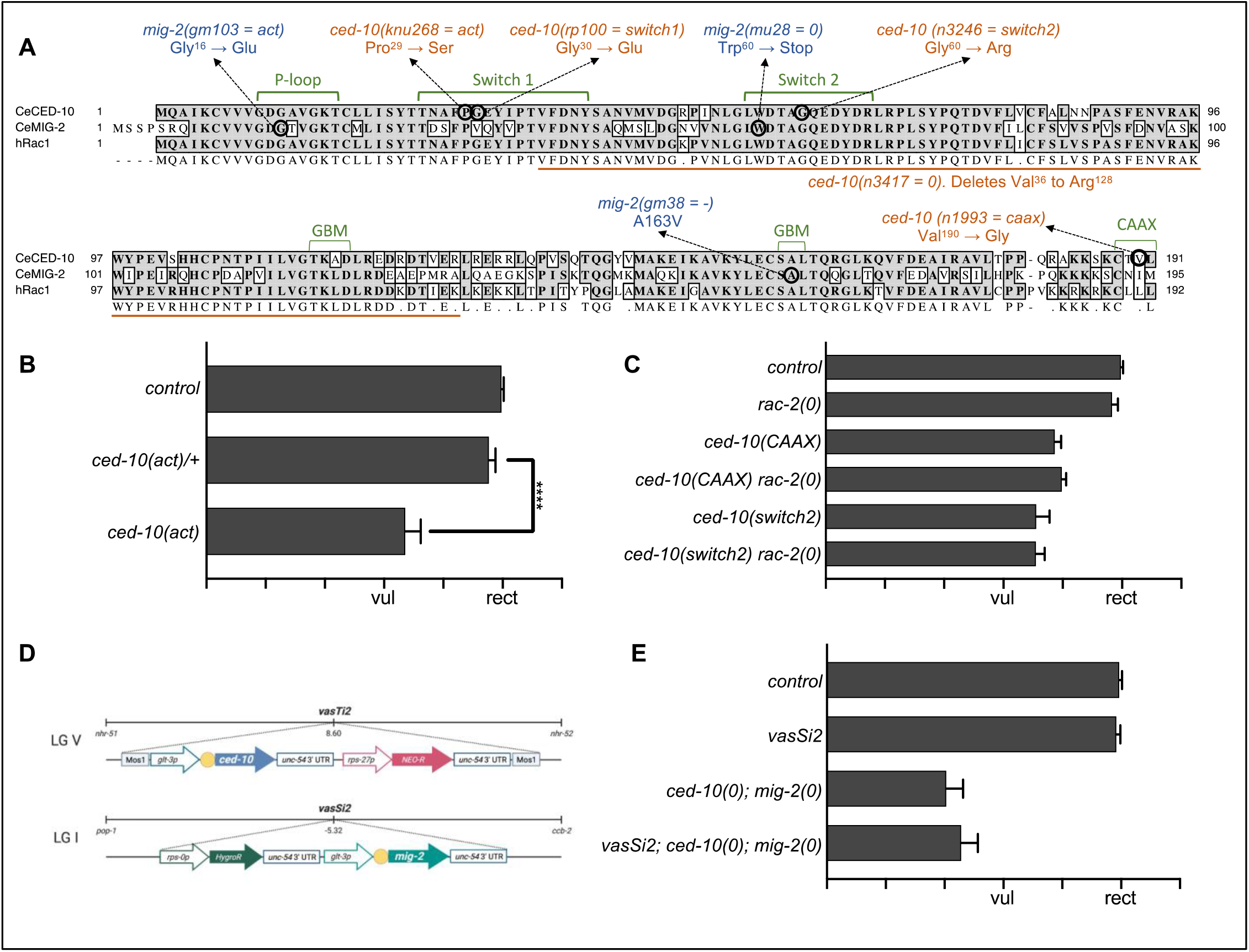
*ced-10/Rac, rac-2/Rac, and mig-2/RhoG mutants and transgenes*. (**A**) Sequence alignment of CED-10, MIG-2, and human Rac1. Domains required for GTPase activity and localization are highlighted in green. Mutations in *ced-10/Rac* and *mig-2/RhoG* used in this study are marked to identify the altered amino acid residue and allele name. (**B**) *ced-10(act)* is recessive. (**C**) A deletion of *rac-2/Rac* on its own, or combined with *ced-10/Rac* mutations, does not affect *ExCa* outgrowth compared to controls. (**D**) Illustration of single-copy insertion transgenes *vasTi2* [*glt-3*p*::FLAG::ced-10*] and *vasSi2* [*glt-3*p*::FLAG::mig-2*]. FLAG epitope is shown as yellow circle. (**E**) The transgene *vasSi2* does not rescue *ced-10(0); mig-2(0)* outgrowth defects.

## Notes

### Competing Interest Statement

The authors have declared no competing interest.

